# A nanobody that recognizes a 14-residue peptide epitope in the E2 ubiquitin-conjugating enzyme UBC6e modulates its activity

**DOI:** 10.1101/700609

**Authors:** Jingjing Ling, Ross W. Cheloha, Nicholas McCaul, Zhen-Yu J. Sun, Gerhard Wagner, Hidde L. Ploegh

## Abstract

A substantial fraction of eukaryotic proteins is folded and modified in the endoplasmic reticulum (ER) prior to export and secretion. Proteins that enter the ER but fail to fold correctly must be degraded, mostly in a process termed ER-associated degradation (ERAD). Both protein folding in the ER and ERAD are essential for proper immune function. Several E2 and E3 enzymes localize to the ER and are essential for various aspects of ERAD, but their functions and regulation are incompletely understood. Here we identify and characterize single domain antibody fragments derived from the variable domain of alpaca heavy chain-only antibodies (VHHs or nanobodies) that bind to the ER-localized E2 UBC6e, an enzyme implicated in ERAD. One such VHH, VHH05 recognizes a 14 residue stretch and enhances the rate of E1-catalyzed ubiquitin E2 loading in vitro and interferes with phosphorylation of UBC6e in response to cell stress. Identification of the peptide epitope recognized by VHH05 places it outside the E2 catalytic core, close to the position of activation-induced phosphorylation on Ser184. Our data thus suggests a site involved in allosteric regulation of UBC6e’s activity. This VHH should be useful not only to dissect the participation of UBC6e in ERAD and in response to cell stress, but also as a high affinity epitope tag-specific reagent of more general utility.

**Highlights:**

- Identified single domain antibodies (VHHs) that bind to UBC6e with high affinity
- VHH binds to a short linear epitope near a phosphorylation site
- VHH binding to UBC6e enhances enzymatic activity
- VHH binding inhibits UBC6e phosphorylation in cells

## INTRODUCTION

Most secretory proteins in mammalian cells fold and acquire modifications in the endoplasmic reticulum (ER) prior to export and secretion. Proteins unable to reach their proper folded state, despite multiple attempts at chaperone-assisted folding, must be degraded. Many misfolded proteins and unfolded proteins are transported or dislocated from the ER back into the cytosol for degradation by the proteasome. This process is termed ER-associated degradation (ERAD). ERAD involves many components, including sensor proteins that recognize the unfolded substrates, the dislocon that facilitates removal of substrates from the ER for delivery to the cytosol, the ubiquitination machinery that poly-ubiquitinates substrates for degradation, as well as the proteasome that degrades the substrates. Collectively, ERAD may comprise several distinct pathways, each with its unique substrate preference and composition of component parts. Understanding ERAD is essential for interpreting the biology of immune cells with high secretory burdens, such as activated B cells and plasma cells. ERAD may contribute to antigen cross-presentation as well.

Hrd1 is an E3 ligase that acts as part of the complex that translocates substrates from the ER lumen to the cytosol: auto-ubiquitination of Hrd1 triggers this translocation^1^. A cryo-EM structure of *Saccharomyces cerevisiae* Hrd1 in complex with its adaptor protein Hrd3 shows that Hrd1 forms an aqueous channel embedded in the lipid membrane^2^. Hrd1 also contains a RING-H2 domain, which specifies its E3 activity^3^. E3 enzymes collaborate with E2 ubiquitin-conjugating enzymes for catalysis. Human Hrd1 has shown *in vitro* ubiquitination activity in conjunction with the E2 UBC7 (UBE2G2)^4^. The E2 UBC6e, also known as UBE2J1, interacts with Hrd1^5^, and was the predominant E2 co-immunoprecipitated with the Hrd1-SEL1L complex^6^, suggesting that UBC6e is the primary E2 that serves Hrd1. Paradoxically, elimination of UBC6e by genetic means leads to accumulation of several functional ER chaperones known to target folding intermediates for ERAD, and therefore can accelerate ERAD^7^. UBC6e thus fine-tunes ERAD activity. We therefore sought to better understand how the activity of UBC6e is controlled.

UBC6e is one of two ER tail-anchored E2s that are the mammalian homologs of the *Saccharomyces cerevisiae* E2 UBC6p^8, 9^. Both homologs, UBC6 (UBE2J2) and UBC6e participate in ERAD, as expression of catalytically inactive mutants exerts a dominant-negative effect and delays degradation of ERAD substrates^9^. The ER membrane tail anchor is essential for proper function of UBC6e: overexpression of a mutant that lacks the transmembrane domain fails to restore the function of UBC6e in UBC6e^−/−^ cells^7^. Provision of a heterologous transmembrane segment only partially restores activity^7^. UBC6e is cleaved by Enterovirus 71 3C^pro^ in its C-terminal region for release from the ER. This cleavage causes a disturbance of normal ERAD in the course of viral infection^10^. Genetic ablation of UBC6e is not lethal in mice but causes sterility in males homozygous for the deletion^11^. UBC6e is phosphorylated at Ser184 in response to cell stress, a reaction thought to be catalyzed by PERK^12^, whereas MK-2 is the kinase responsible for phosphorylation of UBC6e upon inhibition of protein synthesis^13^.The significance of UBC6e phosphorylation remains unclear, but it has been suggested that Ser184 phosphorylation affects both the rate of ubiquitin (Ub) loading onto UBC6e as well as the rate at which UBC6e itself is degraded by the proteasome^12, 14^.

Here we developed an immunological tool to modulate activity of UBC6e. Fragments that correspond to the variable region of heavy chain-only antibodies (VHHs), also known as nanobodies, are the binding domain of heavy-chain only antibodies found in camelids. They retain many of the properties of conventional antibodies, such as high affinity and specificity for their targets. They range from 10-15 kDa in size and can be produced recombinantly in the cytosol or periplasm of *E. coli* in high yield^15^. Many VHHs do not require glycosylation or disulfide bond formation to retain their proper fold and can therefore bind to their targets when expressed in the cytosol of mammalian cells. We have used cytoplasmically expressed VHHs to block nuclear import of nuclear protein (NP) during influenza infection^16^, to protect cells from VSV infection^17^, as well as to track the formation of Asc filaments during inflammasome activation^18^. We generated VHHs that target endogenous UBC6e as a new tool to explore its activity. The use of one such VHH, VHH05, identifies a segment of UBC6e outside of the predicted catalytic center, yet critical for its activity. What makes VHH05 particularly useful as well is its ability to recognize a 14-residue peptide epitope in Ubc6e. It is one of the rare VHHs capable of recognizing such an epitope tag with high affinity and where the epitope tag is readily transferable to heterologous proteins.

## RESULTS

### Identification of VHHs that specifically recognize UBC6e

We immunized an alpaca with a purified recombinant fragment of UBC6e (residues1-234) to evoke an antibody response. UBC6e(1-234) and UBC6e(1-197) were produced using a previously described protocol^19^. A strong antibody response directed against UBC6e was observed after immunization as inferred from an immunoblot prepared with serum from the immunized animal (Supporting figure 1a). While this experiment does not distinguish between heavy chain-only and conventional antibodies, we can nonetheless use the mere presence of antibodies as a proxy for successful immunization. The VHH-encoding regions were selectively amplified from a lymphocyte cDNA library and inserted into phagemid plasmids. After two rounds of panning, 7 VHHs with distinct complementarity determining regions (CDRs) were identified by sequence analysis (Supporting figure 1b). Because CDR1 and CDR2 are germ-line encoded and differed for the newly identified VHHs, we also conclude that these 7 VHHs were all derived from unique VDJ recombination events. Among all VHHs identified, VHH05 exhibited the tightest target binding (low nM dissociation constant). We therefore chose to focus on this VHH for further experiments (Table 1). VHH05 effectively immunoprecipitated recombinant UBC6e but not UBC6 (41% identical in residues 1-147, weak homology thereafter), the ER-associated E2 UBC7, nor any of the other E2s tested (Figure 1a, Supporting figure 1c). We then functionalized VHHs equipped at the C-terminus with a sortase-recognized LPETGG sequence using G_3_-biotin and previously developed sortagging methods^20^. We used the resulting site-specifically biotinylated VHHs for immunoblotting. In contrast to many VHHs that only recognize their targets in their native conformation such as VHH enhancer, which recognizes native but not unfolded GFP, VHH05 detects UBC6e also after denaturation in SDS (Figure 1b)^21^.

**Table 1.** Dissociation constants for the 7 unique VHHs identified. Values represent mean ± standard deviation. Binding was measured using Octet as described in the method section.

| Kd(nM)<br>VHH | UBC6e (1-234) | UBC6e (1-197) |
| --- | --- | --- |
| UBC6e 02 | 80 ± 20 | 60 ± 30 |
| UBC6e 03 | 160 ± 40 | 280 ± 60 |
| UBC6e 05 | 5 ± 2 | 6 ± 2 |
| UBC6e 08 | 900 ± 200 | 600 ± 300 |
| UBC6e 11 | 2200 ± 1500 | 4600 ± 1300 |
| UBC6e 16 | 800 ± 200 | No Binding |
| UBC6e 18 | 1000 ± 700 | 1500 ± 600 |

**Figure 1.**
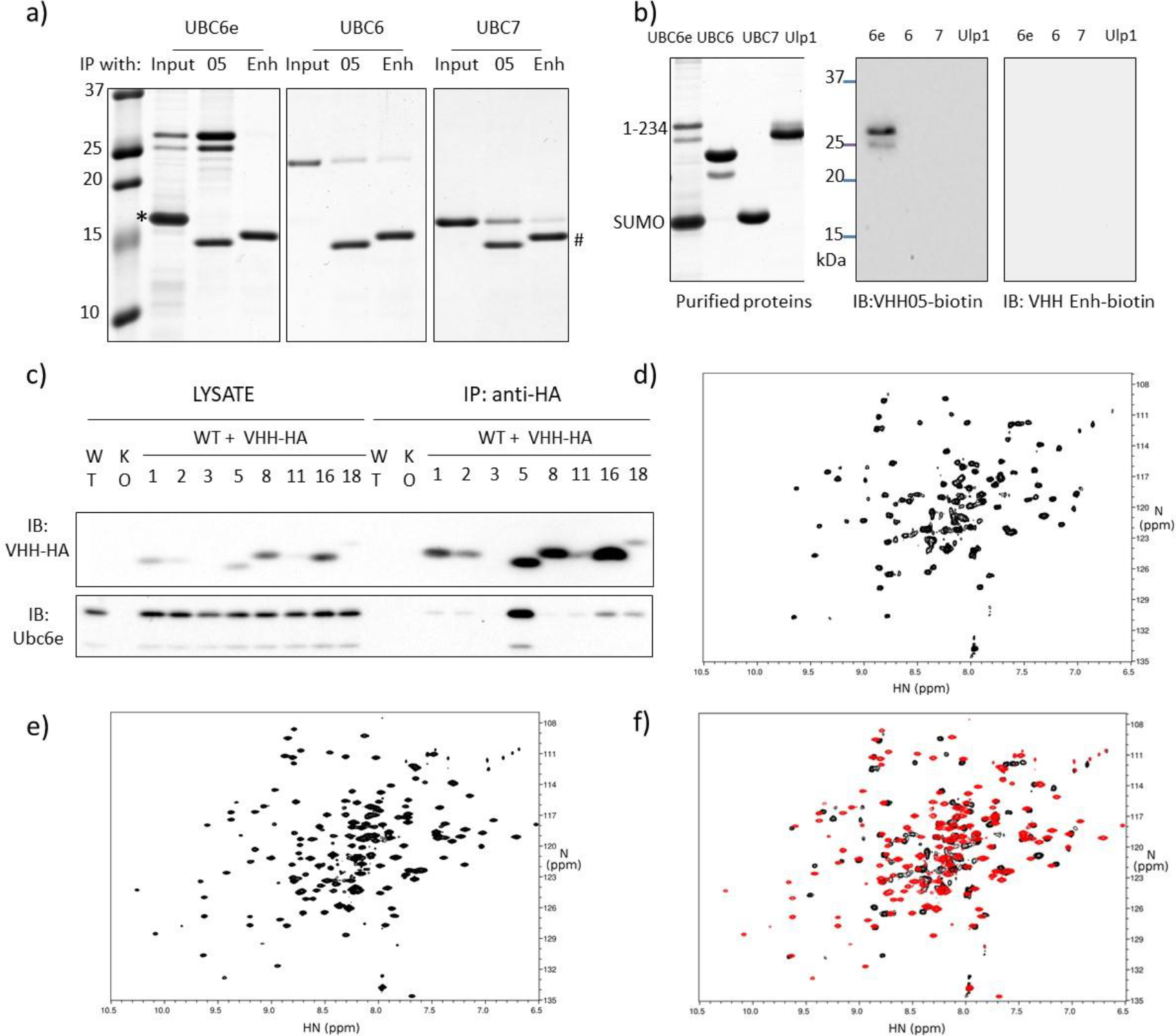
VHH05 can specifically recognize UBC6e with high affinity and good specificity. (**a**) VHH05 can specifically immunoprecipitate UBC6e. Recombinant UBC6e(1-234), UBC6(1-197) and UBC7 were incubated with sepharose bead-conjugated VHH 6E05 or VHH enhancer that is specific to GFP. *: excess SUMO. #: VHH05 and VHH enhancer. (b) VHH05 can specifically immunoblot for UBC6e. Ulp1: SUMO protease. (c) Endogenous UBC6e can be co-immunoprecipitated with VHH05-HA expressed in MEF cells. WT and KO lanes are samples from MEF cells not transfected with VHH. Numbers identify the VHH construct transduced into the cell. (d-f) Addition of VHH05 induces structural changes in UBC6e. HN-TROSY-HSQC spectrum of 0.2mM ^15^N-labeled UBC6e (1-197) in the (d) absence or (e) presence of 0.22 mM unlabeled VHH05. (f) Overlay of data in panels d (black) and e (red).

In contrast to conventional antibodies, many VHHs do not require intrachain disulfide bonds to fold and maintain their function in the reducing environment of the cytosol^16–18^. To test whether the VHHs we identified could be expressed and bind their target in the mammalian cytosol, we expressed the indicated VHHs, equipped with HA tags, in mouse embryonic fibroblasts (MEFs) and performed immunoprecipitations with anti-HA conjugated beads. Endogenous UBC6e was recovered along with VHH05-HA (Figure 1c). None of the other VHHs enriched for UBC6e by immunoprecipitation, possibly because of their comparatively lower affinities. Moreover, not all of the VHHs were expressed at levels similar to VHH05. We did not pursue VHHs besides VHH05 further.

We attempted to characterize the interaction between VHH05 and UBC6e using methods of structural biology. Although VHHs can serve as crystallization chaperones, our numerous attempts at obtaining crystals of UBC6e(1-197) alone or in complex with VHH05 were unsuccessful. Efforts to crystallize UBC6e(1-165) and UBC6e(1-234) in the presence of VHH05 were also unsuccessful. Previous crystallization screening studies also failed to provide a crystal structure of UBC6e^22^. In the absence of a crystal structure, we used NMR to characterize the interaction of UBC6e and VHH05. We produced highly purified ^15^N labeled UBC6e(1-197) for analysis by NMR. For ^15^N-labeled UBC6e(1-197) in solution, we observed disperse peaks consistent with a folded protein mixture, but further efforts failed to arrive at useful structural information (Figure 1d-f). Upon inclusion of unlabeled VHH05, we noticed a major improvement in the quality of the spectra obtained. While still insufficient to derive structural details, this experiment nonetheless suggests that the presence of VHH05 in complex with UBC6e(1-197) broadly alters the structural properties of the E2, consistent with the imposition of a greater degree of order on the protein in solution.

### VHH05 accelerates the loading of UBC6e with Ub and E3-catalyzed UBC6e-Ub hydrolysis

E2 enzymes participate in multiple steps within the ubiquitination cascade. Ubiquitin-activating enzyme E1 activates the C-terminus of free ubiquitin to produce a mixed anhydride intermediate and then forms a thioester between ubiquitin and the E1’s active site cysteine. This ubiquitin is then transferred onto a ubiquitin-conjugating enzyme E2 active-site cysteine, again in thioester linkage. The ubiquitin-loaded E2 can either transfer its ubiquitin in thioester linkage onto a HECT-type E3 ubiquitin ligase for subsequent substrate ubiquitination, or it can be recruited by a RING-type E3 to the substrate and directly transfer the ubiquitin from the E2 onto the substrate(s) (Figure 2a).

**Figure 2.**
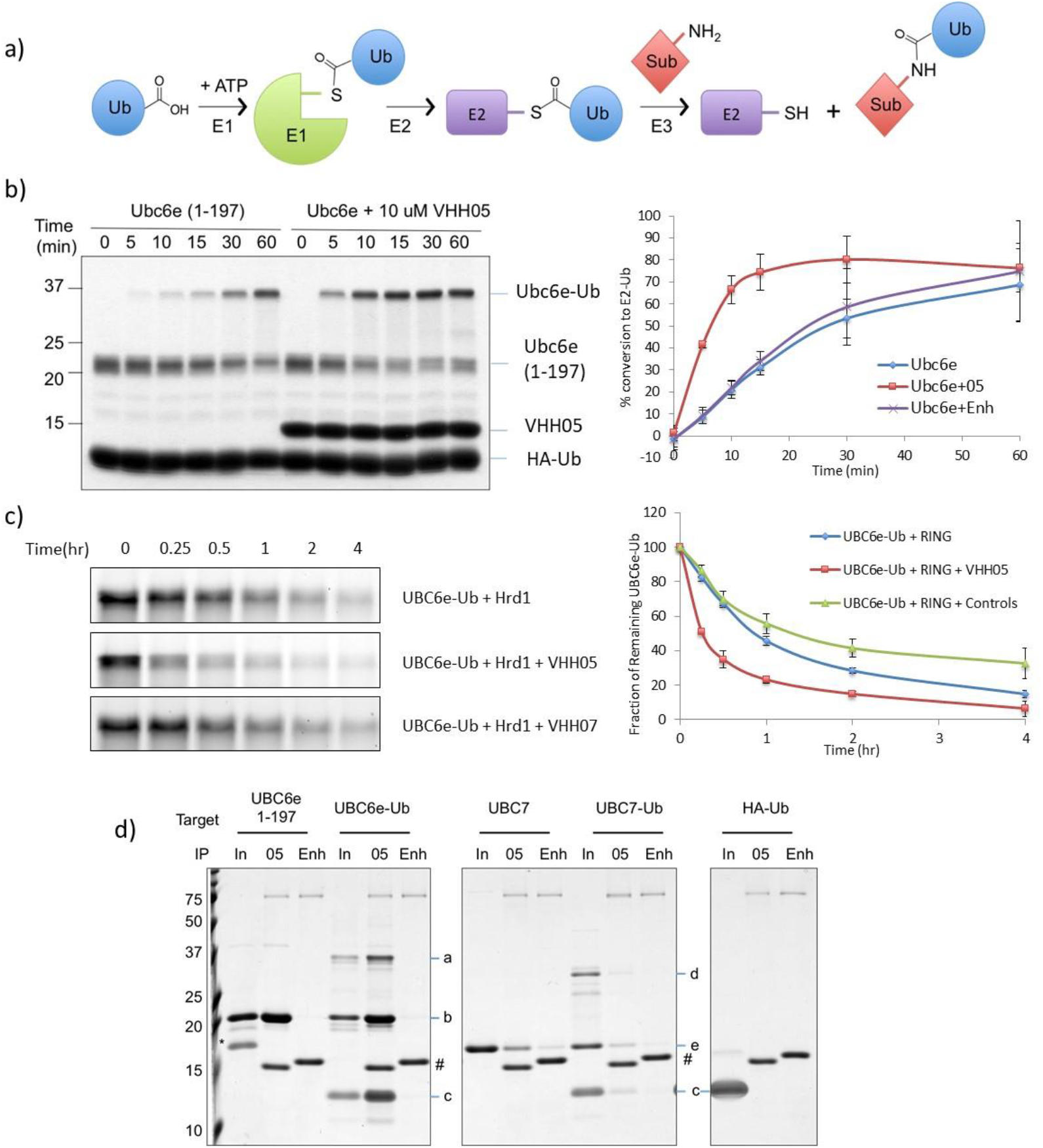
VHH05 accelerates the enzymatic activity of UBC6e. (**a**) Schematic representation of ubiquitin pathway. (**b**) VHH05 accelerates the E2-loading of UBC6e. Ubiquitination reactions were quenched at the indicated time by addition of SDS-containing sample buffer and resolved by SDS-PAGE on a 15% non-reducing gel (left) and quantified by densitometry (right). Data points represent mean ± standard deviation. (**c**) VHH05, but not an MHC-II specific VHH (VHH07), accelerates the hydrolysis of UBC6e-Ub. Samples were prepared and analyzed as in panel b. (**d**) VHH05 binds to both UBC6e and UBC6e-Ub. E2, E2-Ub thioester conjugates or HA-Ub were diluted using tris buffered saline and immunoprecipitated with VHH05- or enhancer-conjugated sepharose beads as described in the methods section. Samples were eluted in 4% SDS with 100 mM and resolved with 15% SDS-PAGE. Since elution was carried out with DTT the thioester has been partially hydrolyzed. For IP labels “in” indicates input; “05” indicates VHH05 immunoprecipitation; “Enh” indicates enhancer immunoprecipitation. a: UBC6e(1-197)-Ub. b: UBC6e(1-197). c: HA-Ub. d: UBC7-UB. e: UBC7. *: SUMO. #: VHH05 and VHH enhancer.

Loading of ubiquitin onto UBC6e can be performed with purified recombinant proteins. UBC6e-Ub thioester is stable in the course of purification by FPLC as well as during analysis by non-reducing SDS PAGE. Because UBC6e(1-234) migrates as two distinct polypeptides on SDS-PAGE, we analyzed a truncated construct, UBC6e(1-197), its design based on the construct used for the crystallization of its homolog UBC6 (pdb: 2F4W)^22^. Both UBC6e(1-197) and (1-234) can be loaded with ubiquitin using recombinant E1 (Figure 2b and Supporting figure 2a). Upon inclusion of VHH05 in the reaction mixture, the rate of loading is accelerated by ~3-fold. This acceleration is observed with the UBC6e-specific VHH05, but not with the GFP-specific VHH, enhancer. VHH05 has no effect on the rate of Ub loading for UBC6e homologs UBC6 or UBC7 (Supporting figure 2b), neither of which VHH05 recognizes.

We then tested whether the presence of VHH05 likewise accelerated the hydrolysis of UBC6e-Ub. The addition of RING domains from E3 enzymes significantly increases the rate of hydrolysis for many E2-Ub thioesters^23^. We therefore used recombinant GST-Hrd1 RING domain (272-343) for this experiment^4^. The molecular weight of the GST-Hrd1 RING fusion protein is very similar to that of UBC6e-Ub, causing these species to overlap on SDS-PAGE. To observe the behavior of UBC6e-Ub without such interference, we expressed UBC6e(1-197) with a C-terminal, sortase-compatible LPSTGG motif and labeled it with Gly_3_-tetramethylrhodamine (TAMRA) using sortagging. We successfully loaded TAMRA-sortagged UBC6e with ubiquitin at rates and with yields similar to those for its unlabeled counterpart (data not shown). In the absence of GST-Hrd1 RING, Ubc6e-Ub hydrolysis proceeded slowly. Addition of VHH05 had little effect on the rate of hydrolysis of UBC6e-Ub itself (Supporting figure 3). On the other hand, addition of GST-Hrd1 RING resulted in substantial UBC6e-Ub hydrolysis within the 4 h time period analyzed, and addition of VHH05 further increased the rate of UBC6e-Ub disappearance (Figure 2c). We confirmed that VHH05 binds to UBC6e-Ub in an immunoprecipitation experiment (Figure 2d). We therefore suggest that the binding of VHH05 to UBC6e promotes a conformation that allows acceleration of both E2 loading and unloading.

### VHH05 binding is independent of phosphorylation of UBC6e

UBC6e can be phosphorylated on S184. This phosphorylation occurs both during ER stress and upon inhibition of protein synthesis^14^. Based on the epitope identified for VHH05, the site of phosphorylation (S184) maps very close to, if not within the binding site of VHH05. We first tested whether VHH05 distinguished between phosphorylated and non-phosphorylated UBC6e. We recombinantly expressed a phosphomimetic mutant (S184E) of UBC6e and tested whether VHH05 still bound. We used VHH05-sepharose to immunoprecipitate UBC6e (S184E), which was recovered as efficiently as UBC6e(1-197) (Supporting Figure 4a). We then randomly biotinylated UBC6e(S184E) on exposed Lys residues using biotin-N-hydroxysuccinimide ester (NHS) and measured its binding to VHH05 using Octet. The K_d_ for the association between VHH05 and UBC6e(S184E) was estimated at 15 ± 10 nM, indistinguishable from that for similarly biotinylated UBC6e(1-197) (14 ± 8 nM) and UBC6e(1-197)-LPETGGG-biotin (16 ± 4 nM), the latter produced by sortagging. In addition, the extent and efficiency of loading of UBC6e(S184E) with Ub is also similar to that of UBC6e(1-197), both in the presence or absence of VHH05 (Supporting Figure 4b). We also tested whether VHH05 immunoprecipitated phosphorylated UBC6e produced in response to cycloheximide (CHX) treatment. VHH05 indeed immunoprecipitates both non-phosphorylated UBC6e and phosphorylated UBC6e (Supporting Figure 4c). We conclude that VHH05 binding is not dependent on the phosphorylation state of its epitope.

### VHH05 inhibits UBC6e phosphorylation in cells

We then tested whether intracellularly expressed VHH05 affects the phosphorylation of UBC6e in response to stress. Cytosolic expression of VHH05 suppressed phosphorylation of UBC6e induced by inhibition of protein synthesis through application of cycloheximide (CHX), imposition of ER stress using dithiothreitol (DTT), or thapsigargin (Tg), as well as by inhibition of E1 using the small molecule inhibitor MN7243 (E1 inhibitor) (Figure 3a). Intracellular expression of a VHH (VHH68), a VHH that targets an unrelated protein, the hemagglutinin protein of influenza virus, had no effect on phosphorylation of UBC6e. Consistent with previous observations, the protein synthesis inhibitor cycloheximide was the most potent inducer of UBC6e phosphorylation tested^13^.

**Figure 3.**
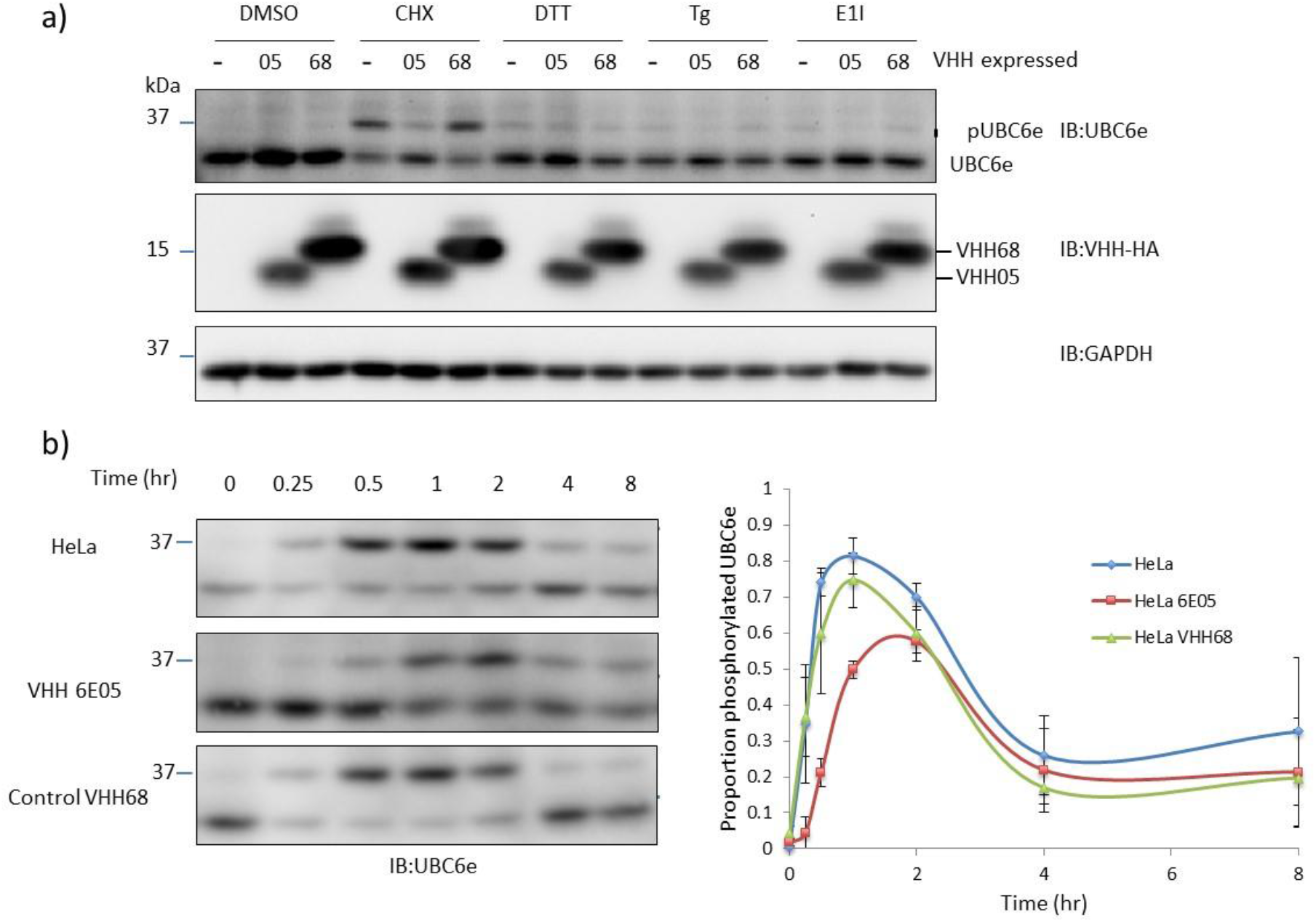
VHH05 blocks phosphorylation of UBC6e when expressed in mammalian cells. (**a**) Phosphorylation of UBC6e in HeLa was reduced by cytosolic expression of VHH05, but not VHH68 that is specific to the nuclear protein of influenza virus. Treatments: 1% DMSO for 30 mins, 50 μM cycloheximide (CHX) for 30 mins, 10 mM DTT for 1 hour, 10 μM Thapsigargin (Tg) for 2 hours, 25 μM MN2743 (E1I, E1 inhibitor) for 1 hour. (b) VHH05 slows down and reduces the phosphorylation of UBC6e when HeLa cells were treated with 50 μM CHX for the time indicated. Quantified data (right) depict mean ± standard deviation and lines serve only to guide the eye.

Cycloheximide treatment induced phosphorylation of ~80% of UBC6e within an hour, followed by slow reversal by cytosolic phosphatases. Cytosolic expression of VHH05 delays peak phosphorylation of UBC6e to 2 hours and decreases the maximum level of phosphorylation to ~60% of UBC6e (Figure 3b).

### VHH05 does not impact the rate of ERAD in live cells

Since VHH05 enhances UBC6e activity (Figure 2) we tested whether ERAD is also accelerated. Using a previously described K562 cell line that stably express the ERAD substrate α1-antitrypsin null Hong Kong variant (NHK) fused with GFP (NHK-GFP), we introduced VHH05 tagged with the fluorescent protein TD-tomato by transient transfection (Supporting Figure 5)^24^. We confirmed that the NHK-GFP was expressed in a doxycycline inducible manner and transfection led to VHH-tomato expression as evaluated by flow cytometry (Supporting Figure 5b). To test whether VHH05 expression impacted the rate of NHK-GFP degradation we performed a cycloheximide pulse-chase experiment. The measurement of GFP levels in doxycycline-treated live cells and those with detectable levels tomato fluorescence (Supporting Figure 5c) revealed that fluorescence decreased over time following cycloheximide chase (Supporting Figure 5d-f); however, VHH05 expression did not substantially alter the rate of NHK-GFP degradation. This observation suggests that UBC6e enzymatic activity is not a primary determinant of the rate of ERAD, at least under these conditions.

### Identification of VHH05 epitope on UBC6e

Since VHH05 can be used to detect SDS-denatured UBC6e by immunoblot, we speculated that it might recognize an extended, less structured segment rather than a 3-dimensional, conformationally constrained epitope. Furthermore, VHH05 recognizes construct UBC6e(1-197) but not UBC6e(1-165), both by immunoprecipitation and by immunoblot (Figure 4a-b). In line with this finding, VHH05 also fails to accelerate loading of UBC6e(1-165) with Ub (Supporting Figure 6). We therefore reasoned that the epitope for VHH05 must reside between residues 166 to 197 in UBC6e. The amino acid sequence for this region is G**<u>S</u>**GSSQADQEAKELARQISFKAEVNSSGKTIA, where the underlined serine residue was mutated to glycine to create a GGG motif to facilitate sortagging of the peptide onto heterologous proteins. Accordingly, the peptide **G<u>G</u>G**SSQADQEAKELARQISFKAEVNSSGKTIA was sortagged onto eGFP-LPSTGG and the resulting chimeric product was then tested for binding to VHH05. eGFP-UBC6e(168-197), but not eGFP-LPSTGG-His_6_, was recovered on VHH05-conjugated sepharose beads (Figure 4c). Further immunoprecipitation experiments demonstrated that VHH05-sepharose retrieves eGFP-UBC6e(168-186) as efficiently as sepharose beads functionalized with VHH Enhancer (VHH_Enh_, anti-GFP) (Figure 4d), indicating that residues 187-197 from UBC6e are not essential for VHH05 binding. VHH05 sepharose, but not Enhancer-sepharose, also recovered the unrelated bacterial protein Arl3 sortagged with the UBC6e(168-186) tag. Biotinylated eGFP-UBC6e(168-197) exhibited a dissociation constant (K_d_ at 7 ± 2 nM) similar to that of UBC6e(1-197) as measured using Octet (Figure 4e), showing that the engrafted region indeed includes the VHH05 epitope. Assessment of biotinylated peptides fragments from this region using Octet showed that VHH05 can bind peptides as short as 14 residues (Kd ~ 0.15 nM).

**Figure 4.**
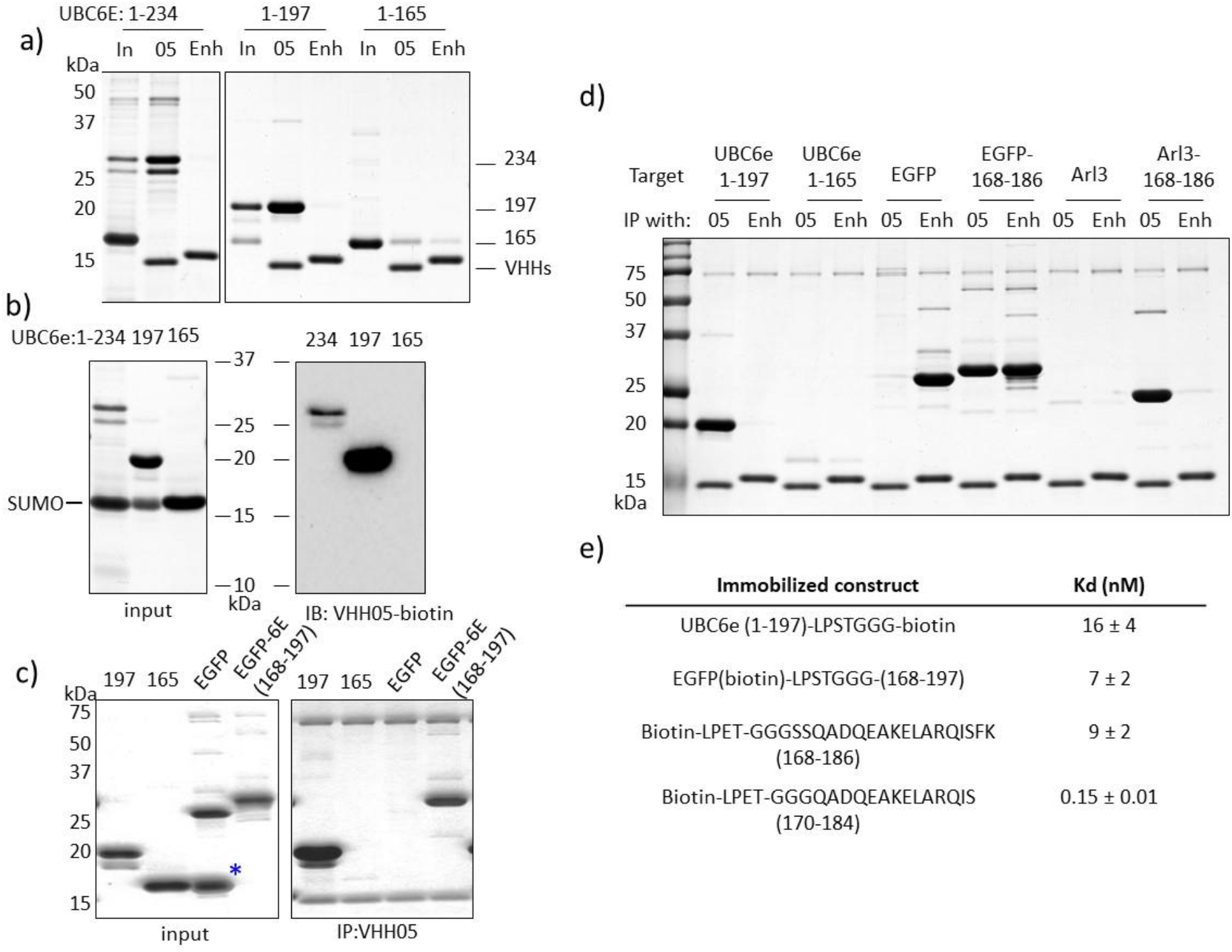
The epitope of VHH05 is located within the UBC6e(168-186) region. (**a**) VHH05 can immunoprecipitate UBC6e(1-234) and (1-197), but not (1-165). “In” indicates input; “05” indicates immunoprecipitation with VHH05-functionalized sepharose beads; “Enh” indicates immunoprecipitation with Enhancer-functionalized sepharose.(**b**) VHH05 does not recognize UBC6e(1-165) on an immunoblot. (**c**) GGG-UBC6e(168-197) was enough to confer binding property to otherwise unrecognized eGFP. (**d**) The helical region (168-186) was sufficient to confer recognition of eGFP and Arl3 by VHH05. (**e)** Summary of binding affinities measured for UBC6e(1-197),epitope tagged eGFP, and biotinylated peptides as measured by Octet.

We performed immunoprecipitation experiments using eGFP functionalized at the C-terminus using sortagging with epitope tags recognized by previously identified VHHs^25–28^ (Figure 5). VHH enhancer (VHH_Enh_) immunoprecipitated all variants of eGFP tested. VHH05 immunoprecipitated eGFP tagged with the 14-mer UBC6e tag as well as VHH_Enh_. The immunoprecipitation by VHH05 was slightly more efficient than that achieved by a VHH that recognizes an epitope tag from beta-catenin (VHH_BC2_) and the correspondingly tagged eGFP^26, 27^. A VHH reported to recognize a 4-residue fragment from α-synuclein (NbSyn2, named VHH_EPEA_ here)^25,28^ failed to immunoprecipitate under these conditions. These experiments unambiguously map the epitope recognized by VHH05 to the segment composed of residues 170-184, with binding independent of the remainder of UBC6e.

**Figure 5:**
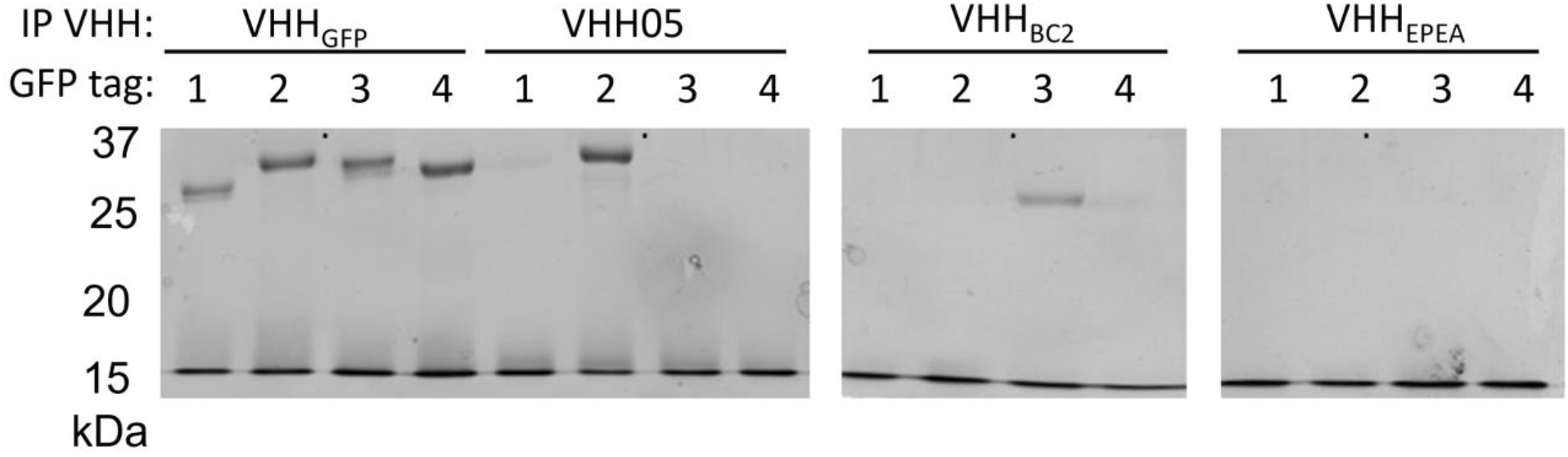
Comparison of immunoprecipitation properties of VHHs that recognize linear epitope tags with Coomassie staining. All VHHs besides VHH05 were previously reported^25–27^. The identity of the GFP tags are: 1- GG (no tag), 2-QADQEAKELARQIS (UBC6e 14-mer), 3-PDRKAAVSHWQQ (BC2 tag), 4- EPEA (EPEA tag). GFP tags were appended to the C-terminus of eGFP using sortagging with identity verified using LC/MS (data not shown). VHHs biotinylated using sortagging were used to immunoprecipitate tagged GFPs. The same preparation of tagged GFP was used for all of the immunoprecipitations with each of the different VHHs. VHHs and GFPs were incubated at equivalent concentrations (5 μM) in TBS + 10% glycerol followed by addition of streptavidin sepharose beads. Beads were washed with TBS + tween 20 and the immunoprecipitate was analyzed using SDS-PAGE on 15% polyacrylamide gels.

## DISCUSSION

ERAD machinery is important to maintain ER function through removal of misfolded proteins before they form aggregates that may be toxic. During ER stress, in which an excess of unfolded protein accumulates in the ER, an unfolded protein response (UPR) is activated through three separate branches: PKR-like ER kinase (PERK), Inositol Requiring Enzyme 1 (IRE1), and Activating Transcription Factor 6 (ATF6)^29–31^. Hrd1 and SEL1L are upregulated during ER stress via the IRE1 and ATF6 pathways, respectively^32–34^. Each of the ER-associated E2s, UBC6, UBC6e and UBC7, is also upregulated during the UPR^12, 35^.

Such transcription-mediated responses may take hours to take effect, yet cells can mount responses to misfolded proteins more rapidly^6^. It would make sense for the cell to activate ERAD as quickly as possible to meet the increasing demand for protein disposal. Our study confirms that UBC6e can respond to cellular stress within an hour as indicated by phosphorylation on Ser184. Further studies are needed to clarify whether S184 is critically important for any aspect of ERAD.

We have generated new VHHs as a tool to identify a region that allosterically regulates the enzymatic activity of UBC6e. VHHs have proven to be particularly useful as reagents for modulating the activity of enzymes^36–38^. VHHs have been identified that inhibit^37,39^ or augment enzymatic activity^39, 40^, through direct active site binding^37, 41^ or via allosteric mechanisms^42,43^. VHHs that enhance enzymatic activity are rare. In the characterization of extensive libraries of VHHs found to bind enzymes, only 3-5% of those identified impart enhancement of enzyme activity^39,43^. VHH05, which binds to a region of UBC6e outside the active site and does so with a dissociation constant in the low nanomolar range, enhances the rate of Ub loading on UBC6e by ~3-fold (Figure 2b). Inclusion of VHH05 also increases the rate of UBC6e-Ub hydrolysis in the presence of the Hrd1 RING domain (Figure 2c). Despite the observation that VHH05 potentiates UBC6e activity *in vitro*, expression of VHH05 did not affect the rate of degradation of the ERAD substrate α1-antitrypsin Null Hong Kong variant (Supporting Figure 6). The NMR binding data support the imposition of a global conformational change in UBC6e when complexed with VHH05 (Figure 1d-f). Although we lack a structure of the catalytic domain of UBC6e at atomic resolution, the observation that UBC6e(1-165) retains activity (Supporting figure 3) indicates that the segment recognized by VHH05 lies outside this core.

Allosteric regulation of activity has been reported for many E2s. Binding of E3 RING domains allosterically enhances E2 reactivity for transfer of Ub onto substrates^44^. As an example, UBCH5b activity is enhanced upon binding to multiple E3s^45, 46^. In line with these precedents, we likewise observed an enhanced rate of UBC6e-Ub hydrolysis upon inclusion of the Hrd1 RING domain (Figure 2c). Non-RING domain-mediated allosteric regulation of E2s has been reported also. UBC7 is activated by a non-RING domain on its cognate E3 ligase gp78 (ref 46, 47). Non-covalent association with ubiquitin can activate the E2 (UBCH5B)-Ub-RING E3 complex^48^. Phosphorylation of “gateway residues” in E2s can open the active site and enhance enzymatic activity. CDC34 (Ub-loading), UBC3B (substrate degradation) and UBE2A (substrate ubiquitination) are all activated by phosphorylation of gateway residues^49–52^. *In vitro*, the phosphomimetic mutant S184E UBC6e is loaded with Ub at a rate similar to that of WT UBC6e (Supporting figure 4), and therefore Ser184 on UBC6e is unlikely to serve as a traditional gateway residue. However, we propose that the larger stretch in UBC6e (168-186) is a “gateway segment” that when bound, improves access to the active site. VHH05 binds to this segment and enhances UBC6e activity. Again, with reference to the NMR data, it is clear that a global conformational alteration results in UBC6e(1-197) from the binding of VHH05. Ser184 phosphorylation could induce association with phosphoserine-binding proteins, and similarly facilitate allosteric enhancement of UBC6e activity. If such phospho-Ser184 interactors exist, their identity remains to be established. Nonetheless, we speculate that interactions of the 1-197 fragment with endogenous cellular proteins might exert a similar effect to regulate UBC6e activity *in vivo*.

This speculation finds support in the impact of cytoplasmic expression of VHH05 on phosphorylation of Ser184. The phosphorylation site Ser184 falls within the segment recognized by VHH05. VHH05 binds to both phosphorylated and non-phosphorylated epitopes, showing that Ser184 is not essential for VHH05 binding. However, cytosolic expression of VHH05 reduced phosphorylation of UBC6e induced by inhibition of protein synthesis. This suggests that VHH05 competes with intracellular regulators or kinases for binding to this target site, confirming that the VHH epitope is adjacent to this Ser184 phosphorylation site. Intracellular expression of VHH05 is thus effective in altering the network of protein-protein interactions in which UBC6e normally participates.

Phosphorylation of UBC6e is induced both by ER stress and by inhibition of protein synthesis^12, 13^. Activation of UBC6e, via phosphorylation, caused by the accumulation of unfolded proteins or the inhibition of protein synthesis, might serve to enhance ERAD in response to severe cellular stress. We found that TLR agonists such as LPS (TLR4), CpG (TLR9), Imiquimod (TLR7), Malp2 (TLR2/6), and Pam3CSK4 (TLR1/2) rapidly (within 20 minutes) induce phosphorylation of UBC6e^53^. Since the kinase MK2 is also activated in TLR-engaged dendritic cells^54, 55^ and activated MK2 induces strong phosphorylation of UBC6e, it is likely that MK2 is also responsible for TLR-induced phosphorylation of UBC6e. Since MK2 is activated in the course of diverse cellular processes, phosphorylation of UBC6e may provide a mechanism by which a state of cellular stress can be conveyed to the ER, where ERAD can be pre-emptively activated in preparation for the impact of cytosolic stress on the ER environment. There is no reason to assume that such relay of information would be unidirectional, exclusively from ER to cytosol.

The tightest binding VHH identified in this study (VHH05) was shown to bind with high affinity to a short (14 amino acid) stretch of UBC6e (Figure 4e, Figure 5). There are few reported VHHs-peptide pairs that engage in high affinity interactions. Notable examples include the NbSyn2-EPEA system from an α-synuclein-binding VHH^25^, the BC2-Spot Tag system from a betacatenin binding VHH^26,27^, and the newly reported Alfa tag system from a VHH raised against a designed α-helical peptide^28^. These VHH-peptide pairs have been used for a variety of applications, ranging from traditional molecular biology approaches such as affinity purifications and immunoprecipitations, to more specialized approaches such as super resolution microscopy and immune cell purification workflows^25–28^. Established systems still present some shortcomings. The NbSyn2 VHH binds a very short epitope (EPEA) but requires epitope placement at the C-terminus and the interaction is of relatively low affinity and specificity^25^. The BC2 nanobody binds a peptide epitope of approximately the same length as VHH05 (12 residues for BC2 vs 14 for VHH05); however, beta-catenin is widely distributed at the cell surface and in the nucleus, potentially complicating some applications^26,27^. The Alfa tag VHH recognizes with very high affinity a 14-mer peptide not found in common organisms; however, this tag remains to be tested in diverse settings^28^. Application of the VHH05-peptide interaction reported here should complement the properties and applications of these existing systems.

There are extensive crosstalk and feedback mechanisms that connect protein synthesis, folding, trafficking, with cellular stress. Identification of new tools that can be used to disrupt processes of interest, and to do so specifically without massively interrupting cell function or by physical elimination of proteins of interest remains an important goal. Inhibition of UBC6e phosphorylation and enhancement of enzymatic function by VHH05 offers a fine-tuned approach for dissecting protein function, ERAD and cellular proteostasis in living cells. VHH05 and the approach disclosed here should be valuable for future studies of ERAD and related processes.

## EXPERIMENTAL PROCEDURES

### Antibodies and reagents

Commercial antibodies for OS-9 (monoclonal rabbit; Abcam), GAPDH (Horseradish peroxidase (HRP)-conjugated monoclonal rabbit; Cell Signaling Technology), HRP-conjugated goat anti-mouse Ig and anti-rabbit antibody (Southern Biotech), HRP-conjugated rabbit anti-goat antibody (Southern Biotech) were used according to supplier’s recommendations. Rabbit anti-UBC6e serum (1:3000) was generated and used as previously described^6^. High sensitivity streptavidin-HRP (Pierce) was used at a 1:10,000 dilution. Cycloheximide and doxycycline (Clontech) and MLN7243 (ChemiTek) were obtained from commercial sources. Protein G Sepharose-4 Fast Flow, Protein A Sepharose 4 Fast Flow and Con A Sepharose 4B were purchased from GE Healthcare.

### Plasmids, protein expression and purification

The pCMV-CDH-EF1-puro UBC6e constructs have been described^7^. VHHs were amplified from a cDNA library generated from alpaca lymphocytes and subcloned into pD vector. VHHs enriched after 2 rounds of panning were subcloned to the pHEN6 vector for expression in E. coli and to pINDUCER20 vector for expression in mammalian cells as described^16–18^.

UBC6e constructs were expressed using the Champion pET SUMO system (Thermo Scientific). Plasmids encoding SUMO-UBC6e(1-234), (1-197) or (1-165) were transformed into LOBSTER BL21(DE3) *E. coli*. His-tagged SUMO-UBC6e was purified by binding to Ni-NTA beads, washed with 10 mM imidazole in PBS, and eluted with 250 mM imidazole in 50 mM Tris, 500 mM NaCl, pH 8.0. 50 μg of SUMO protease was added and the solution was dialyzed into 50 mM Tris, 150 mM NaCl, pH 7.4. Cleaved Ubc6e was purified from SUMO using fresh Ni-NTA beads followed by washing with 20 mM imidazole in 50 mM Tris, 500 mM NaCl. UBC6e was further purified by size exclusion on a S75 column (GE Healthcare, UK). Typical yields were 5-10 mg per 1L culture.

GST-hmHrd1 RING (272-343) was expressed using the pGEX-6P-1 plasmid^4^. Bacterial lysates were collected as described above and the lysates were incubated with 1 ml Glutathione Sepharose 4B (GE Healthcare, UK) at 4°C for 1 hour. The beads were washed 3 times with 10 ml PBS, and GST-Hrd1 RING was eluted with 10 mM reduced glutathione in TBS. The eluted protein was further purified by size exclusion on a S75 FPLC column.

VHHs were expressed using the pHEN6 vector. Plasmids coding for PelB-VHH-LPETGG-His6 were transformed into WK6 *E.coli*. Bacteria were grown on ampicillin selection at 37 °C and protein expression was induced with 1mM IPTG at 30 °C overnight. Bacterial pellets were collected and resuspended in TES buffer (50 mM Tris, 650 μM EDTA, 2 M sucrose, 15 mL buffer per liter of culture) to prepare for osmotic shock. After incubating for 2 hours at 4 °C, 75 mL distilled H2O was added and the bacterial suspension was incubated overnight at 4 °C. VHHs were purified from the supernatant by binding to Ni-NTA beads then further purified by size exclusion on a S75 column. Typical yields were 20-40 mg per 1L culture.

### Cell culture and transient transfection

Wildtype (WT) and UBC6e^−/−^ MEFs have been described^6^. MEFs were maintained in Dulbecco’s modified Eagle’s medium (DME) supplemented with 10% IFS and 0.0007% v/v β-mercaptoethanol (MEF medium) in humidified air containing 5% CO_2_ at 37°C. HeLa cells were maintained in Dulbecco’s modified Eagle’s medium (DME) supplemented with 10% IFS. K562 human myeloma cells (ATCC) were maintained in RPMI 1640 medium supplemented with 2mM L-glutamine, 10% FBS, 200 μg/mL G418 and 4 μg/mL Blasticidin. VHH05-tdTomato fusions were transfected into K562 cells using an Amaxa 4D-Nucleofector according to manufacturer instructions. In these K562 cells NHK-GFP was by addition doxycycline as previously described^24^. Doxycycline was added approximately 1 h post nucleofection.

### Immobilization of VHHs to CNBr-activated sepharose beads

CNBr-activated sepharose beads (200 mg, Sigma-Aldrich) were washed twice with 20 mL of 1 mM HCl, and neutralized with 100 mM NaHCO_3_, 500 mM NaCl, pH 8.3. 2 mg of each VHH (VHH6E05, VHH enhancer or VHH MHCII-07) in PBS was added to beads in 20 mL of 100 mM NaHCO3, 500 mM NaCl, pH 8.3 and reacted overnight at 4 °C. The beads were blocked with 50 mM Tris, 500 mM NaCl, pH 8.0 for 2 hours at room temperature, and then TBS containing 1% BSA overnight before use.

### Immunoblotting and immunoprecipitation (purified protein)

Unless otherwise stated, 30 ng (as estimated from absorbance at 280 nm) of each protein was loaded for immunoblot. VHH-biotin was used at 1 μg/ml in 5% non-fat milk in Tris-buffered saline containing 0.05% Tween 20. High sensitivity HRP-Conjugated Streptavidin (Pierce, MA) was used at a 1:10,000 dilution.

For immunoprecipitation using immobilized VHH05 and VHH enhancer, 50 μL of beads (corresponding to 100 μg VHH) and 50 μg of target protein was used. Immunoprecipitations were conducted in TBS containing 1% BSA. Beads were washed 3 times before elution in 2x sample buffer (4% SDS, 100 mM DTT).

### Nuclear magnetic resonance analysis of UBC6e

^15^N-labeled UBC6e(1-197) was express from *E.coli* in M9 minimal medium with ^15^NH_4_Cl as sole nitrogen source and purified as described above. ^15^N-TROSY-HSQC spectra were acquired with 0.2 mM ^15^N-labeled UBC6e(1-197), 20 mM NaPO_4_, pH 7.5, with 50 mM NaCl and 10 mM DTT, 10% D_2_O in the absence or presence of 0.22 mM unlabeled VHH05. Data were acquired on Bruker Advance III 800 MHz spectrometer at 25° C. Data were collected with 16 scans per FID, 1024 complex points in direct ^1^H dimension, and 50 complex points in indirect ^15^N dimension for UBC6e alone, and 256 scans per FID, 1024 complex points in direct ^1^H dimension, and 116 complex points in indirect ^15^N dimension for UBC6e with VHH05. The NMR data were processed and visualized using NMRPipe/NMRDraw^56^.

### Generation of stable producer cell lines using lentiviral transduction

Lentivirus was produced in HEK293 cells. HEK293 cells were plated in a 6-well plate and grown until ~70% confluency, and transfected with 1.5 μg of plasmids containing the gene of interest (backbones in pINDUCER20 or pCDH-CMV-MCS-EF1-Puro), 0.65 μg psPAX2 and 0.35 μg pMD2.G. Media containing lipofectamine 2000/plasmids complex were removed after 6 hours of transfection, and HEK cells were grown in DMEM containing 10% heat-inactivated FBS for 2 days. Virus particles were harvested by spinning down the HEK cell pellet and passing the media through a 0.45 μm filter. The media containing virus particles were mixed 1:1 with fresh media and used to transduce MEF or HeLa by incubating for 6 hours. The cells were grown in regular media overnight and selected with 500 μg/mL geneticin for pINDUCER20-transduced cells and 2 μg/ml puromycin for pCDH-CMV-MCS-EF1-Puro-transduced cells.

### Immunoblotting and immunoprecipitation (lysates)

Cultured cells were lysed in HEPES-buffered saline (HBS) with 1% NP-40 supplemented with protease inhibitor cocktail (complete protease inhibitor cocktail tablets; Roche Applied Science) on ice for 20 mins. Cell debris were removed by spinning at 13,000 rpm for 20 min at 4°C. Protein content was determined using a BCA protein assay. Proteins were denatured in sample buffer 2% SDS and 0.1 M DTT, resolved by SDS-PAGE, transferred to PVDF membranes and immunoblotted using standard procedures.

All immunoprecipitation procedures were performed at on ice. Cell lysates were collected in HBS with 1% NP-40 and incubated with CNBr-activated sepharose beads conjugated with VHH05 (50 μg bead volume, 30 μg VHH) or anti-HA agarose beads (Thermo Scientific) overnight. The beads were washed for 3 times with TBS with 1% NP-40 and resuspended in SDS-PAGE sample buffer with 0.1 M DTT.

### Peptide synthesis and purification

Peptides were synthesized using flow-based solid-phase peptide synthesizer using Fmoc chemistry as previously described^57^. Peptides were conjugated to Rink-amide linker resin to produce C-terminal amides. Following synthesis beads were air-dried and peptides were cleaved from resin and deprotected in 92.5% trifluoroacetic acid, 5% H_2_O, and 2.5% TIPS. Peptides were precipitated into cold diethyl ether, air-dried, and purified using reversed-phase C18 HPLC using 10-50% Acetonitrile/H_2_O gradient.

### Sortase reactions

VHH-LPETGG-His6,eGFP-LPSTGG-His6, or peptide-LPETGG preparations incubated with 500 μM of GGG-peptides in the presence of 5 μM heptamutant sortase-His_6_ at room temperature for 1 hour or 4°C overnight. For reactions with VHHs or eGFP, unreacted substrates and sortase were removed by incubating with Ni-NTA beads, and excess peptides were removed using PD10 column. For peptide reactions products were purified by HPLC. A typical yield is 50% after purification.

### E2 loading reactions and UBC6e-Ub hydrolysis reactions

For analytical E2-loading reactions, 3 μM of purified E2 (UBC6e, UBC6 or UBC7) was incubated with 10 μM HA-UB and 100 nM purified His6-mouse E1 in 100 mM Tris with 2 mM ATP, 5 mM MgCl_2_ and 200 μM DTT at 37°C, in the presence or absence of 10 μM VHHs. Reactions were quenched at different time points with equal volume of 2x SDS sample buffer with no reducing agents. Samples were analyzed with non-reducing 15% SDS-PAGE gels.

For the preparation of UBC6e(TAMRA)-Ub, 4 mg of SUMO-UBC6e-LPSTGG in 2 ml TBS was SUMO cleaved with 100 μg of SUMO protease, followed by exposure to 5 μM Heptamutant sortase-His_6_ and 500 μM of G_3_-TAMRA. The sortase reaction mixture was directly used for E2-loading reaction without purification. To the reaction mixture, 10 μM HA-UB and 100 nM purified His6-mouse E1 in 100 mM Tris with 2 mM ATP, 5 mM MgCl2 and 200 μM DTT was added, and the E2 loading reaction was allowed to proceed at 37°C for 45 minutes. The reaction mixture was centrifuged to remove insoluble debris, and directly loaded onto an S75 column for purification. The peak corresponding to UBC6e(TMR)-Ub was pooled and concentrated. The product contains about 50% of unloaded UBC6e (TAMRA), both from incomplete purification and subsequent hydrolysis during purification. The yield was about 20%.

To analyze the hydrolysis of UBC6e(TAMRA)-Ub, 2 μM UBC6e(TAMRA)-Ub was incubated with 10 μM GST-HRD1 RING(272-343) and VHHs (10 μM) in TBS. Reactions were quenched with equal volume of 2x SDS sample buffer with no reducing agents at indicated time points, and analyzed using 15% non-reducing SDS-PAGE. The gel was imaged using absorption at 546 nm for visualization of TAMRA-labeled UBC6e-Ub and UBC6e (ChemiDoc, BioRad).

### Octet measurements

Experiments were performed using the Octet RED96 Bio-Layer Interferometry. The target of interest is biotinylated either through sortagging or labeled with (+)-Biotin N-hydroxysuccinimide ester (Sigma-Aldrich). For NHS labeling, 2-fold molar excess of biotin-NHS was used for labeling at room temperature for 1 hour in PBS. Excess biotin-NHS was removed using ZeBa columns (7K MWCO, Thermo Fisher). All octet measurements were taken in 1% BSA in PBS with 1% Tween. 10 ng/mL of biotin-labeled target were loaded on Dip and Read™ Streptavidin (SA) Biosensors (Pall Fortebio) until the reading exceeded 1 nm. These sensors were then dipped into wells containing different concentrations of VHH05, ranging from 50 nM to 8 μM. Average and standard deviations were taken for at least 3 measurements.

## ACKNOWLEDGEMENTS

We acknowledge the A*STAR NSS PhD fellowship (JJL) and the Cancer Research Institute Irvington postdoctoral fellowship (RWC) for funding. We also acknowledge the Biophysics Instrumentation Facility at MIT for use of the Octet Bio-Layer Interferometry System (NIH S10 OD016326). We acknowledge NIH (5R01AI087879-10, HLP; GM047467 and AI037581, GW) for financial support. We acknowledge Professor Ron Kopito for provision of the K562 cell line expressing GFP-NHK fusions.

