## Supporting Figures for "A nanobody that recognizes a 14-residue peptide epitope in the E2 ubiquitin-conjugating enzyme UBC6e modulates its activity"

Running title: *Modulation of UBC6e function by a nanobody*

[§]Current address: Department of Cancer Biology, Dana Farber Cancer Institute, Boston, MA 02215.

*To whom correspondence should be addressed: Hidde L. Ploegh: Program in Cellular and Molecular Medicine Boston Children's Hospital, Boston, MA 02115; Tel. 617-919-1613

[#]These authors contributed equally to this manuscript

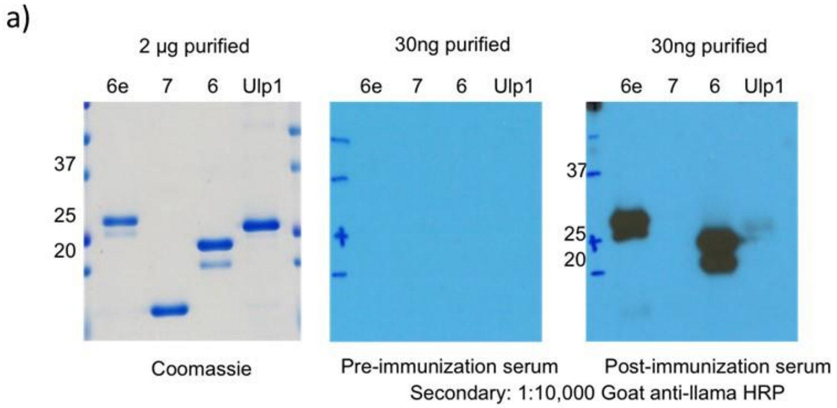


b)

| VHH ID | CDR3 sequence |
|---|---|
| 6E02 | VYYCNANR-GV----TRLRGQGTQVT |
| 6E03 | VYYCHAVRMVG---VLDYWGQGTQVT |
| 6E05 | VYYCSKSG--------AYWGQGTQVT |
| 6E08 | VYYCNADGLGRTGIRAHYWGQGTQVT |
| 6E11 | IYYCNAVRAP----LFNYWGQGIQVT |
| 6E16 | VYYCNAELPG----RSVYWGQGTQVT |
| 6E18 | VYVCNADFAPA---YLKYWGQGTQVT |

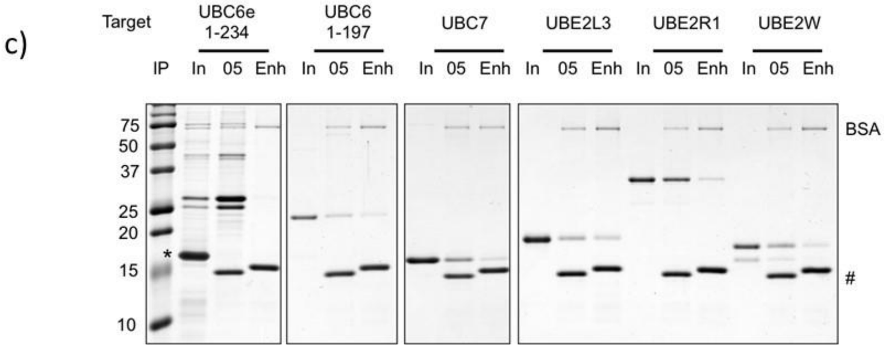


Supporting figure 1. VHH05 is specific to UBC6e. (**a**) The post-immunization serum from alpaca injected with recombinant UBC6e(1-234), UBC7 and UBC6 (1-197) can immunoblot for UBC6e and UBC6, indicating that an antibody response has been elicited in the animal. (**b**) Sequences of Ubc6e specific VHHs CDR3 regions identified by phage display. (c) Among 6 purified recombinant E2s, VHH05 selectively enriches for UBC6e. The immunoprecipitation was performed as described in the methods section. E2s were expressed and purified as previously described[1].

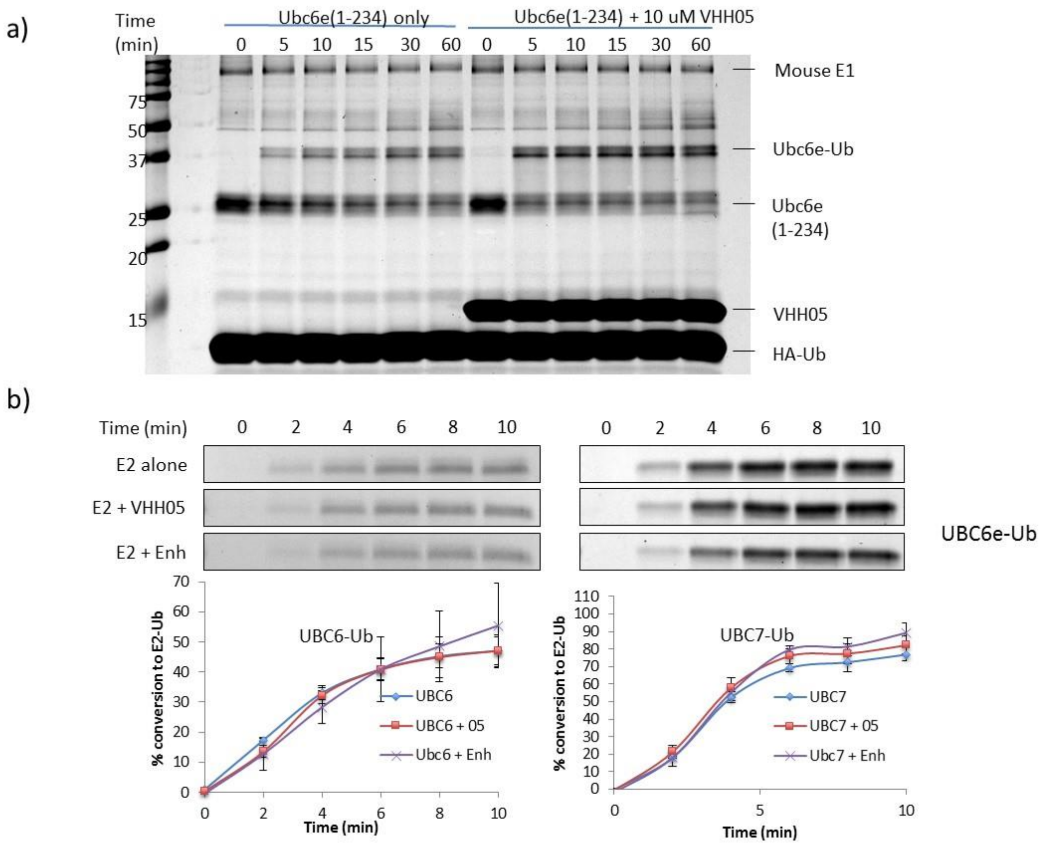


Supporting figure 2. VHH05 specifically accelerates the E2-loading of UBC6e. (a) The loading of UBC6e(1-234) is accelerated in the presence of VHH05. (b) The rate of Ub loading for UBC6 and UBC7 is not affected by VHH05. Quantified data points represent mean ± standard deviation.

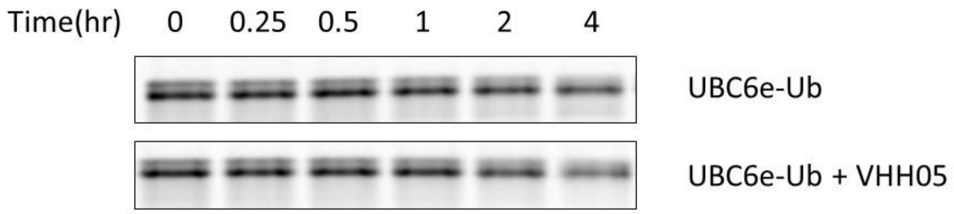


Supporting figure 3. VHH05 does not accelerate the hydrolysis of UBC6e-Ub in the absence of Hrd1 ring domain. Ubiquitination reactions were quenched at the indicated time by addition of SDS-containing sample buffer and resolved by SDS-PAGE on a 15% non-reducing gel.

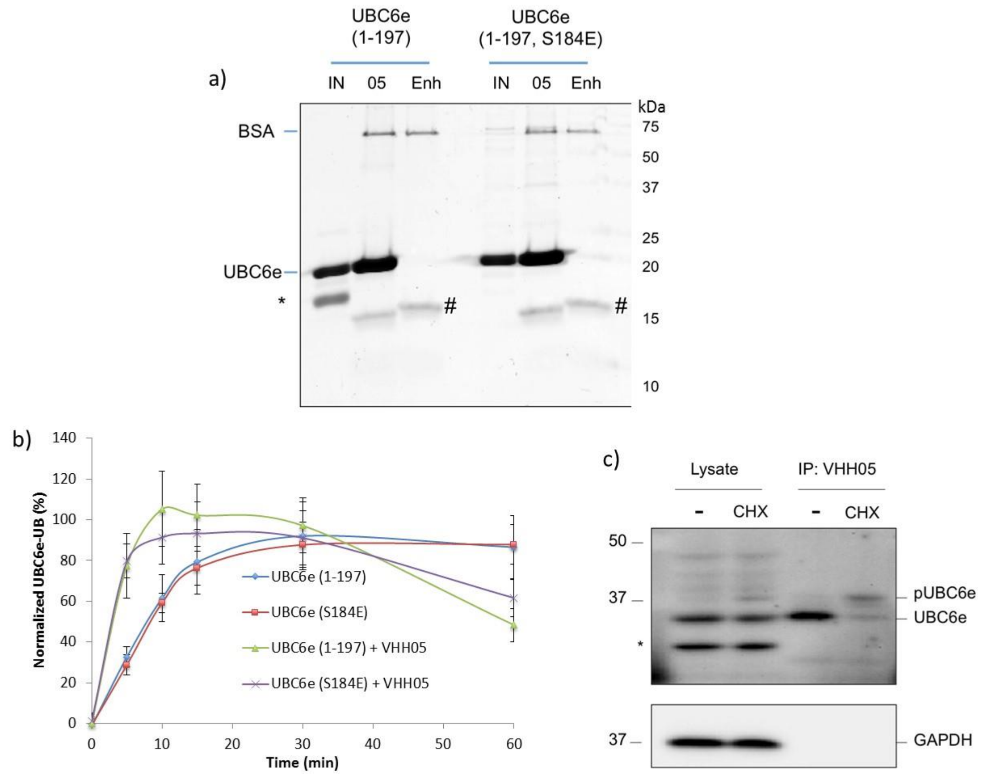


Supporting Figure 4. (**a**) Immobilized VHH05 immunoprecipitates both UBC6e (1-197) and UBC6e (1-197, S184E). *: cleaved SUMO protein from expression as a SUMO fusion. #: VHH05 and VHH enhancer. BSA: Bovine Serum Albumin used for blocking. (**b**) The E2 loading of phosphomimic mutant UBC6e (1-197, S184E) is similar to that of wildtype UBC6e(1-197). The experiment and quantification was performed as described in the methods section. Error Bars represent standard deviation (n=3). (**c**) Immobilized VHH05 immunoprecipitates both UBC6e and phosphorylated UBC6e. HeLa cells were treated with 50 μM cyclohexamide for 30 minutes before collection in 1% NP40. Lysate and IP samples were immunoblotted with 1:3000 polyclonal rabbit anti-UBC6e serum. pUBC6e: phosphorylated UBC6e.

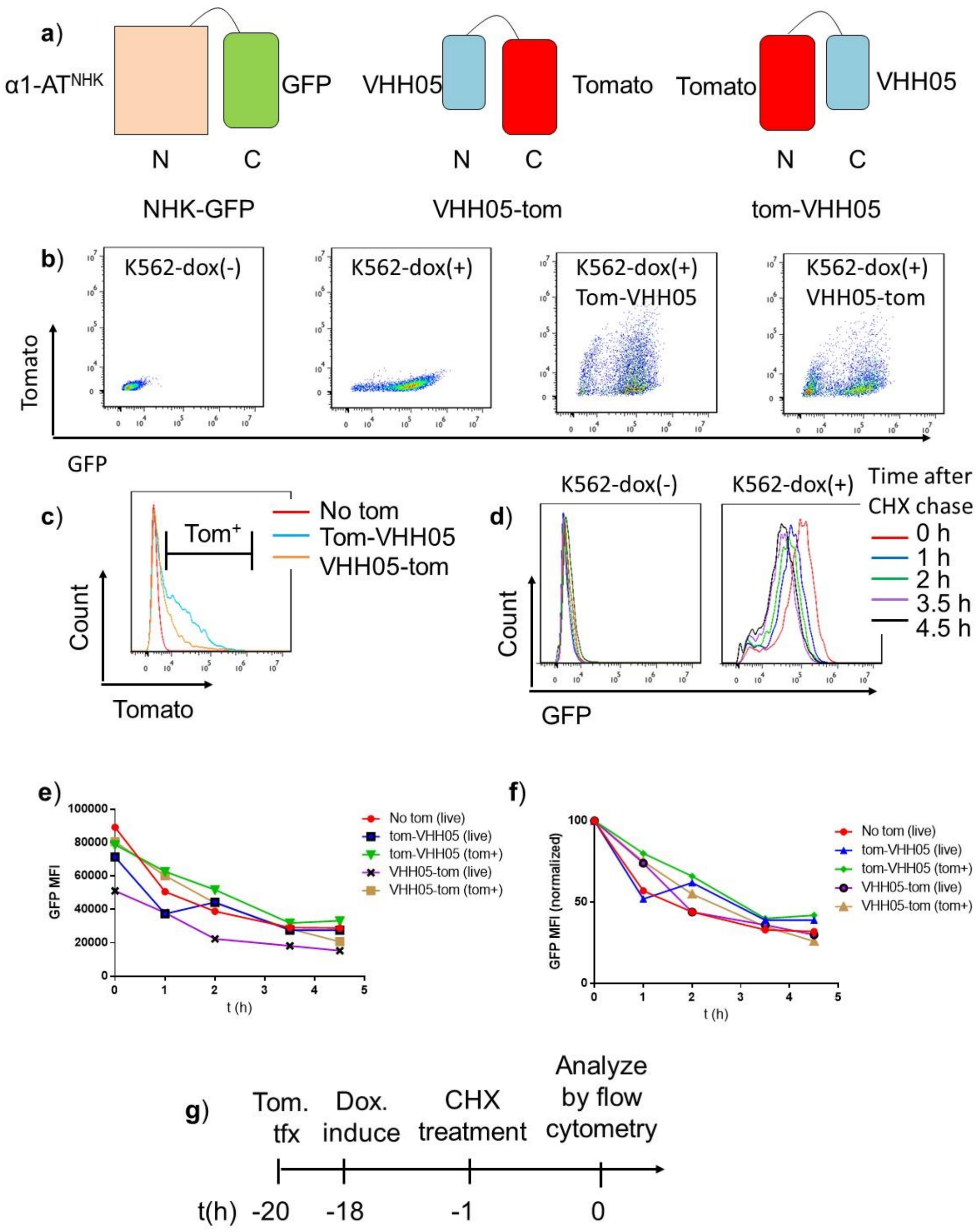


Supporting Figure 5. Characterization of VHH05 expression on the degradation of an ERAD substrate. (**a**) Summary of the protein constructs used. K562 cells that stably express $\alpha$1-AT$^{NHK}$-GFP under a doxycycline inducible promoter were used as previously described[2]. VHH05-TDtomato fusions were applied as described in methods. (**b**) Scatter plots of stably transfected K562 cells with or without doxycycline induction and following transfection with VHH05-tomato constructs. Analysis was performed 18-20 hours after transfection and doxycycline induction. (**c**) sample histrogram showing gating strategy used to analyze tomato positive K562 cells. (**d**) Histograms of GFP expression levels over time in K562 cells either induced with doxycycline [dox(+)] or not induced [dox(-)]. The timeline for the experiment is depicted in panel g of this figure. (**e**-**f**) Plots of the median fluorescence intensity (MFI) for GFP in K562 cells over time following cycloheximide treatment. Values are either raw MFI (**e**) or the MFI for each condition normalized to t = 0. These experiments were performed three times with similar results (data not shown). (**f**). Either live cells (live), as identified by forward and side scatter profile, or tomato positive cells (tom+), as identified as described in panel c, were analyzed. (**g**) Schematic of the timeline for the experimental protocol.

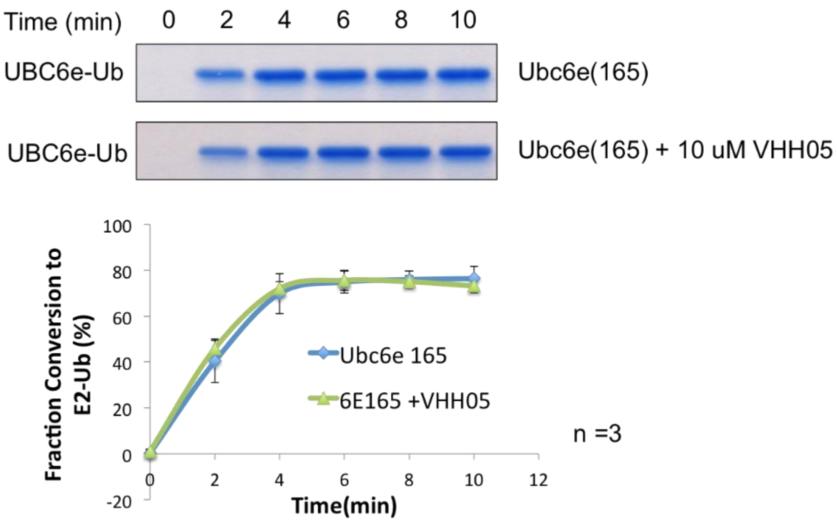


Supporting Figure 6. VHH05 does not affect the rate of formation of UBC6e(1-165)-Ub. The experiment was performed and analyzed as described in the methods section.


1. Jin, J. P.; Li, X.; Gygi, S. P.; Harper, J. W., Dual E1 activation systems for ubiquitin differentially regulate E2 enzyme charging. *Nature* **2007,** *447* (7148), 1135-U17.

2. Leto, D. E., Morgens, D. W., Zhang, L., Walczak, C. P., Elias, J. E., Bassik, M. C., Kopito, R. R., (2018) Genome-wide CRISPR Analysis Identifies Substrate-Specific Conjugation Modules in ER-Associated Degradation. *Mol. Cell 73*, 1-13.
